# Extracellular matrix particle treatment induces digit regeneration in soft-tissue preserved amputation (SPA)model of adult mice

**DOI:** 10.64898/2026.08.05.742898

**Authors:** Yingqin Liu, Boyu Li, Chunmei Bao, Luting Zeng, Zijian Wang, Xin Sun, Guangchen Sun

## Abstract

**Background:** In mouse classic amputation model, second phalanx (P2) is incapable of regeneration. Extracellular matrix (ECM) solution has shown limited ability to induce digit regeneration in classical amputation model of the murine digit. However, the effect of solid ECM particles on bone regeneration is not well understood due to difficulties in treating solid particles in classical amputation model.

**Methods:** We examined the regenerative effects of ECM particles in mice digit by establishing a soft tissue preserved amputation (SPA) model on P2, through removing the amputated bone whilst preserving soft tissue. ECM particles implanted into the amputation site and wrapped in the preserved soft tissues. Bone regeneration was assessed by morphological examination and micro-CT scans.

**Results:** We observed bone regeneration in the SPA model; specifically, new bone formed at the P2 distal end. Implantation of ECM particles exerted a pro-regenerative effect, characterized by increased bone volume and decreased bone density. Moreover, the ECM induced the formation of free-floating bone, further supporting its role in bone regeneration. Combined treatment with ECM particles and bone morphogenetic protein 2 (BMP2) resulted in a significant increase in bone volume.

**Conclusions:** We demonstrate that soft tissue preservation at the amputation site can overcome the intrinsic regenerative limitationsl. Using SPA model, we found that ECM particles have a proven ability to promote bone regeneration, and that the combination of ECM particles with BMP2 further enhances bone regeneration. These findings underscore the therapeutic promise of ECM-based strategies, for clinical translation in non-regenerative finger injuries.

**Summary statement:** Solid-state Extracellular matrix can induce mice digit regeneration and has the potential for clinical application.

**Highlights:** The SPA model we developed enables ECM particles to adhere to wounds, and our research has found that:

1. Digit bone regeneration was shown in SPA model .
2. ECM particles treatment promoted bone regeneration and can generate free-floating bone in SPA model.
3. Combination of ECM + BMP2 treatment induced strong digit regeneration in SPA model.

## 1. Introduction

Amphibians such as salamanders can regenerate their limbs/digits after amputation [1, 2], while mammals have weak regeneration ability, limited to seasonal growth of antlers, closure of ear holes in rabbits/mice, and regeneration of finger/digit tips in humans, monkeys, and rats[3-8], The limited regeneration in the mammals listed above has been achieved by blastema, and such regeneration has been referred to as epimorphic regeneration[9-12]. P3 amputation in mice digits can accomplish a regenerative response through the formation of the blastema. Blastema, a transient population of progenitor cells that form from the blending of periosteal and endosteal/marrow compartmentalized cells that undergo differentiation to restore the amputated structures[13].However, regeneration of P3 is limited to the distal portion of the amputation, and proximal amputation exhibits a progressively weaker regenerative response until the regenerative response disappears[3, 13-16]. In general, the less the P3 is amputated, the stronger the regenerative response becomes.

In human fingertip amputation, despite the absence of bone regeneration, the soft tissues of the fingertip can be restored to relative normalcy, and a new fingerprint could seamlessly rejoin the original fingerprint line without scarring [17, 18]. Such healing of an amputated fingertip without bone tissue lengthening was also referred as regeneration. In 2007, Han et al. proposed that bone lengthening should be the indicator of whether there is digit regeneration, and restoration of soft tissue without bones is considered healing, but not digit regeneration [19]. P2 classical amputation (CA) is the amputation of mouse P2 together with soft tissue in the same plane, which removes surrounding soft tissue. The main feature of healing process of CA is the generation of large amount of cartilage on the lateral side of the periosteum, which forms a chondrogenic callus that encircles the stump, followed by formation of new bone through endochondral ossification. The P2 healing results in an increase in diameter but not in length [20]. According to the definition made by Han et al., the second phalanx(P2) respond to amputation by bone healing in CA, but not bone regeneration [20, 21].

Extracellular matrix (ECM) is predominantly derived from the decellularization of animal organs, including the bladder and small intestine, while advancements in technology have facilitated the production of ECM from *in vitro* cultured cells and synthetic sources. As a biological scaffold material, ECM has been used in the reconstruction of a variety of tissues/organs[22-26]. Studies of ECM used in murine digit amputation-regeneration studies have shown that ECM has effects such as recruitment of cells, enhancement of gene expression, and formation of bone nodules, but did not show the ability to increase bone length or bone volume. These results indicate that ECM may not be able to induce regeneration of mouse digits in CA model. However, the ECM in these studies was in liquid form[27-31]. It is unclear whether the application of ECM particles can induce enhanced bone regeneration.

Other regenerative treatments for P2 such as matrix metalloproteinase 1, electrical stimulation, were unable to induce significant bone regeneration other than transplanted tissues, such as nail bed[29, 32-36]. The use of Bone Morphogenetic Protein (BMP) is an exception as Ide et al. found that neonatal mouse ulna and radius regeneration could be successfully induced using BMP2/BMP7[35, 37]. Furthermore, Yu et al. induced regeneration of mouse digits/limbs with BMP2/BMP7, resulting in a significant increase in bone length[38, 39]. These studies have shown that BMP-induced regeneration is preceded by chondrogenesis and followed by osteogenesis in the form of endochondral ossification, which is the same as osteogenesis in digit/limb development, but different from epimorphic regeneration of P3 via blastema, making BMP2 one of the most promising approaches to induce finger/limb regeneration.

In this study, we established SPA model at P2 and observed bone regeneration in the absence of ECM particles. Treatment of ECM particles induced significant bone regeneration in SPA model. Furthermore, combinational ECM and BMP2 treatments enhanced regenerative effect of ECM particles in a BMP2 dose dependent manner. Our results showed that presence of surrounding soft tissue is critical for regeneration of digit amputation and ECM/ECM+BMP2 are promising targets for development of novel treatment to achieve finger/limb regeneration.

## 2. Materials and Methods

### ECM particles preparation

ECM was prepared as described previously [40, 41]. The basement membrane and underlying lamina propria were isolated and harvested from fresh pork bladder. Next, the membranes were treated with Tris buffer (10-20 mM, pH 8.0), 1% TritonX-100, Tris buffer (pH 7.6) with 1 U/ml RNAse and 50 U/ml DNAse, and deionized H2O, after which they were lyophilized to obtain sheets. The sheets were then comminuted and stored at -80°C. Prior to implantation, the ECM powder was placed in a white pipette tip (10 µl) and manually extruded into particles using a flat-tip needle.

### Animals

Adult Kunming female mice (6-8 weeks) of specific pathogen free grade were purchased from Hunan SJA Laboratory Animal Co. The 2nd and 4th mouse digits (corresponding to the index and ring fingers in humans) of the hindfoot were amputated and the rest digits remained intact. ECM particles were implanted after the amputation in the ECM group, and there was no treatment after the amputation in the control group. Each group included three digits in experiments displayed in Fig.2, 3 and 4 and five digits per group in experiments displayed in Fig.1. The experimental protocol was approved by the Experimental Bioethics Committee of Guilin Medical University. Approval number: GLMC202303159. To avoid systematic errors, we classified the ‘ring finger of the left foot’ and the ‘index finger of the right foot’ as one group, and the rest as the other group. This ensured that both ‘ring and index fingers’ were present in the same group.

**Figure 1.**
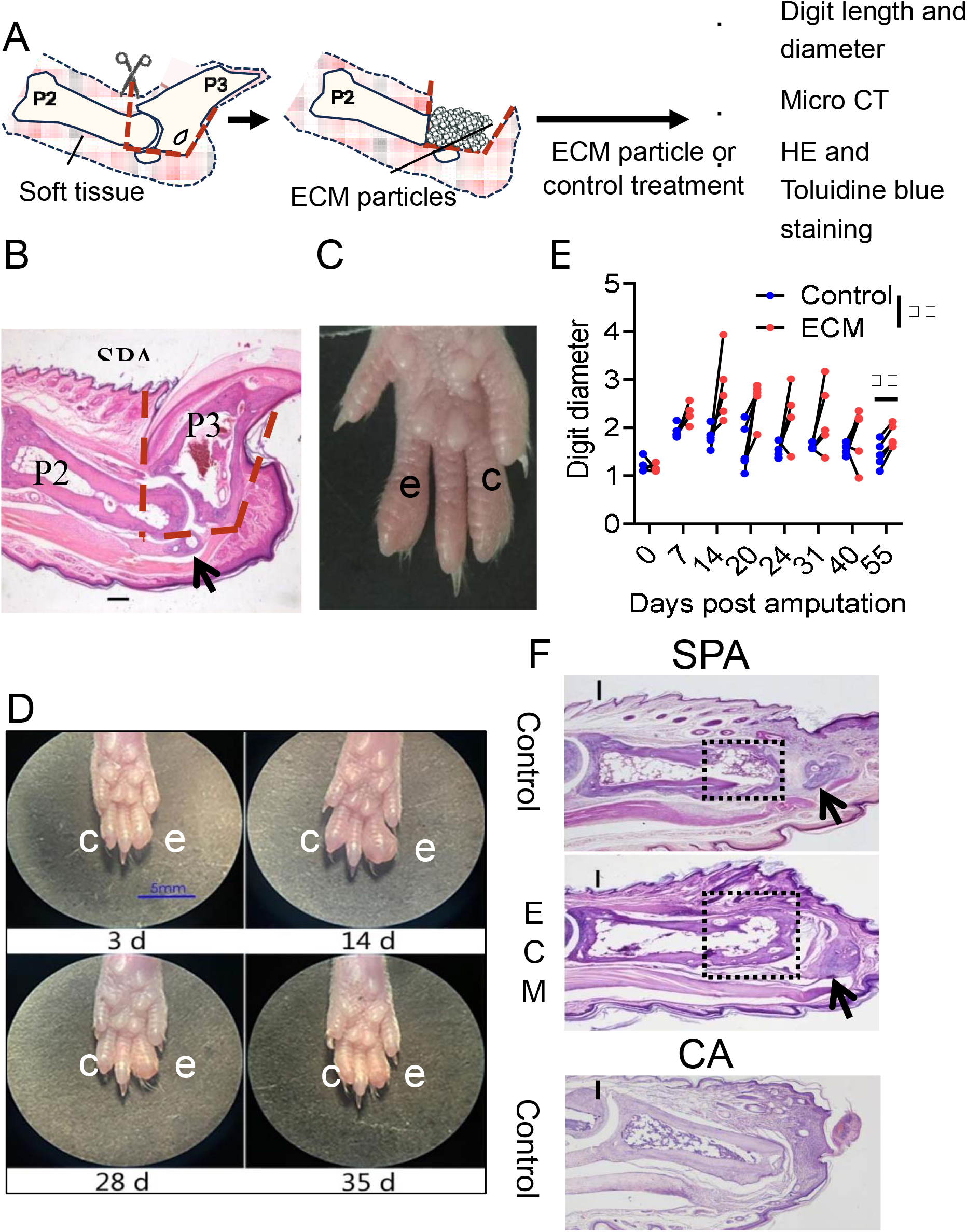
Bone regeneration in the SPA model. (A, B) Schematic and HE staining illustration of establishment of SPA model in mice digit followed by ECM particles treatment. Amputation was performed following the red dashed line to remove P3 and part of P2 while preserving the adjacent soft tissue and sesamoid bone. ECM particles were treated in the site of surgery. (C) SPA model was established on index and ring digits. (D) Appearance of mice digits in SPA models from 3 to 35 DPA. (E) Digit diameter (mm) was measured and compared between control and ECM groups from 0 to 55 DPA. Measurements from paired digits were connected by lines. (F) HE staining of mice digit in CA model with control treatment, SPA model with control treatment, and SPA model with ECM treatment at 40 DPA. Dashed box indicates regenerated bone. 2 way ANOVA with sidak’s multiple comparisons test was used for comparison between ECM and control groups to evaluate the overall land time point specific effect of ECM treatment. n = 5, n indicates the number of biological replicates. **p?0.01. Top or left of images indicates proximal site; Scale bar, 200 µm in (B, F) and 5 mm in (D); Black arrow, sesamoid bone; e, ECM group; c, Control group.

**Figure 2.**
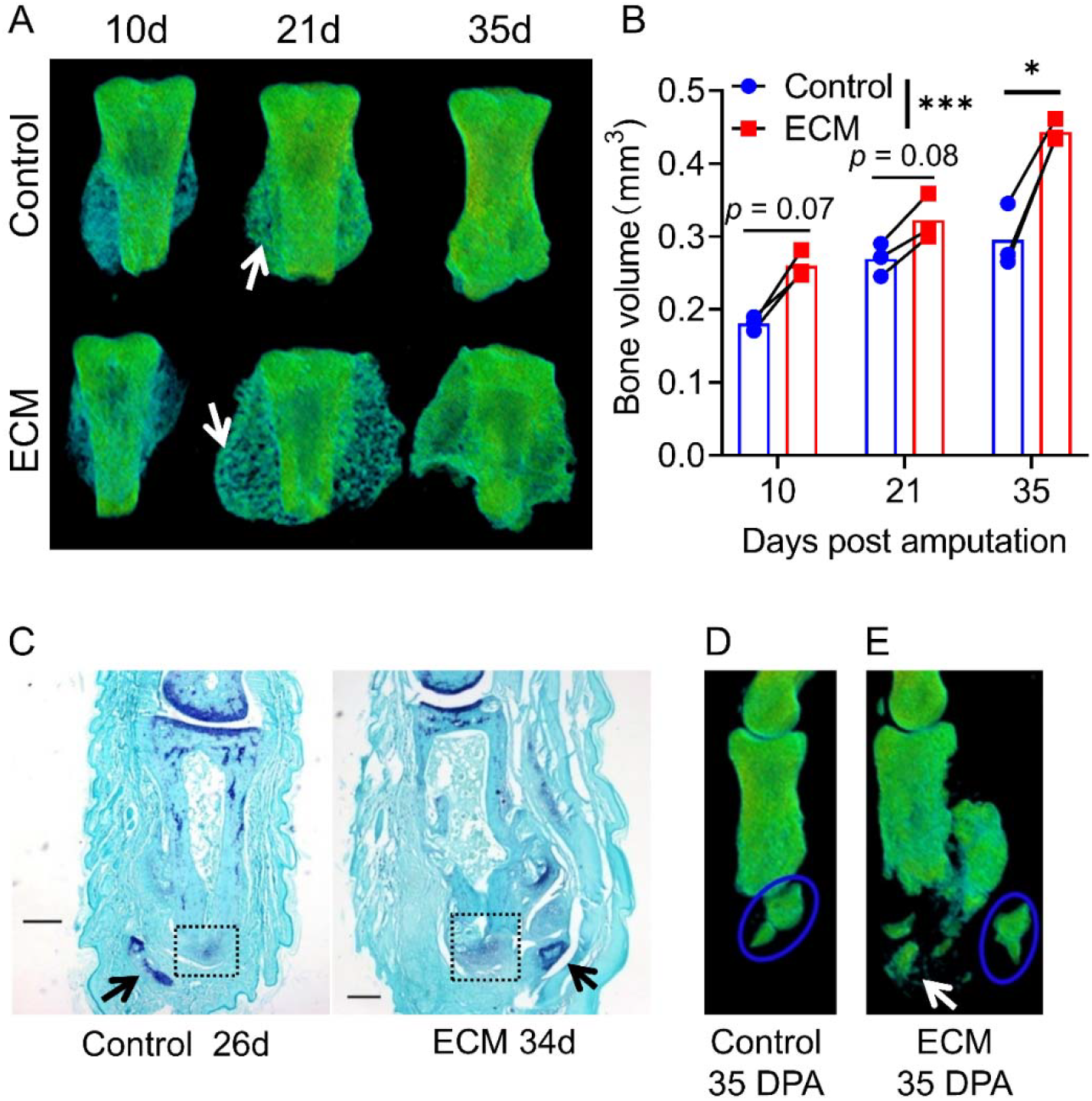
ECM particle treatment promotes generation of new bone in SPA model. (A) micro-CT 3D reconstruction of mice digit in control and ECM groups of SPA model at 10, 21 and 35 DPA. White arrow indicates new bone. (B) Bone volume values of 10, 21and 35 DPA were obtained from micro-CT scans and compared between control and ECM groups in SPA model. Measurements from paired digits were connected by lines.(C) Toluidine blue staining of paraffin sections. The black box shows the cartilage. Black arrow indicates sesamoid bone. (D-E) micro-CT 3D reconstruction of mice digit in (D) control and (E) ECM groups of SPA model at 35 DPA. The blue circle indicates sesamoid bone, white arrow indicates free-floating bone. n = 3, n indicates the number of biological replicates. 2way ANOVA with sidak’s multiple comparisons test was used for comparison between ECM and control groups to evaluate the overall land time point specific effect of ECM treatment. *p < 0.05, ***p < 0.001. Top of images indicates proximal site, and scale bar indicates length of 200 µm in (C).

**Figure 3.**
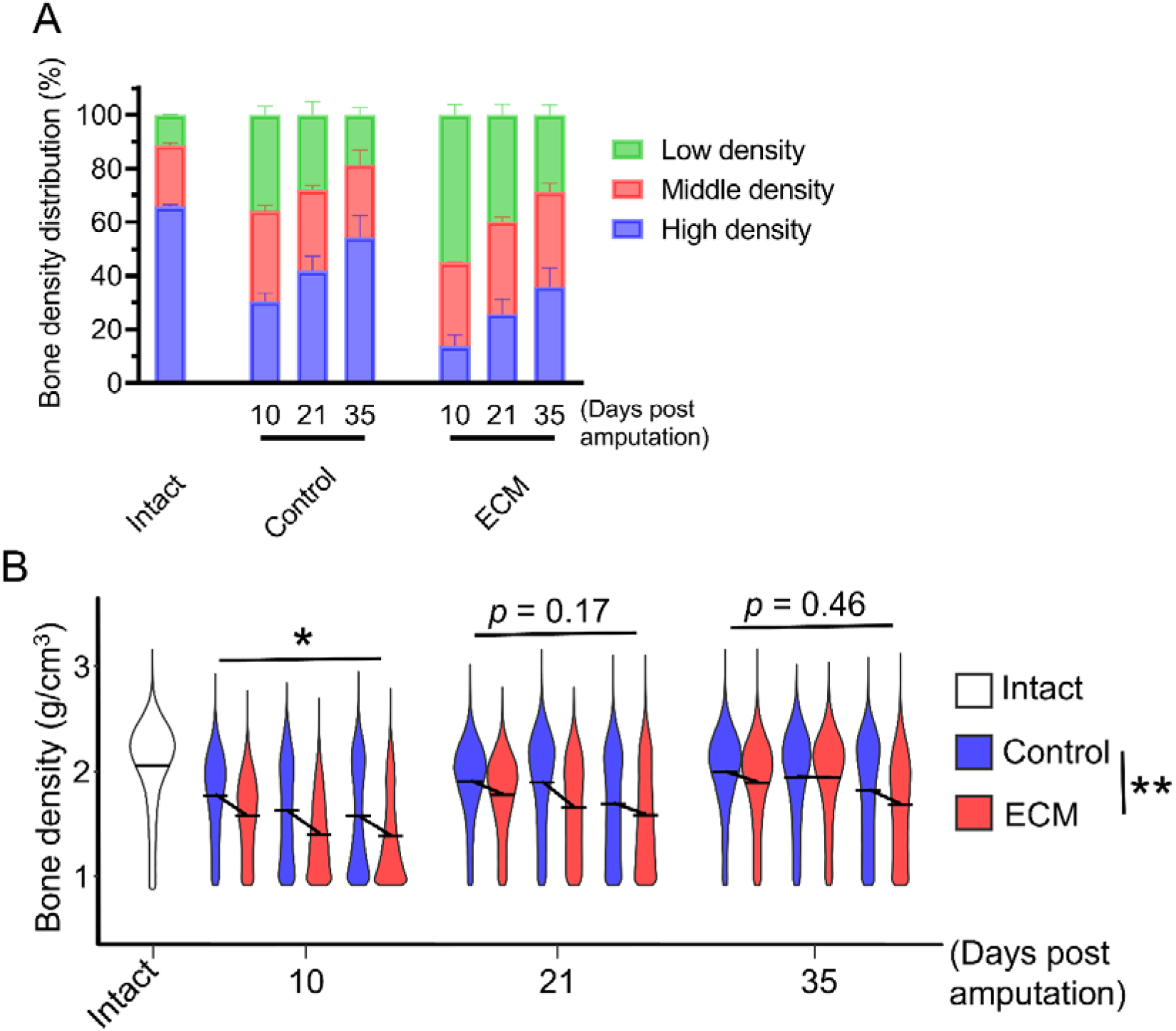
ECM treatment reduces the bone density of P2 in SPA model. (A) SPA model was established in mice digits and received ECM particles and control treatment. Bone densities in intact, control, and ECM groups were obtained by micro-CT scanning and proportion of high, medium, and low bone density zones were calculated at 10, 21, 35 DPA. (B) Mean bone mineral density of P2 were calculated and compared between digits in intact, control, and ECM groups of SPA model from 10 to 35 DPA. Each violin represents measurements from single digits, and mean bone density from paired digits were connected by lines. n = 3, n indicates the number of biological replicates. 2way ANOVA with sidak’s multiple comparisons test was used for comparison between ECM and control groups to evaluate the overall and time point specific effect of ECM treatment. *p< 0.05, **p< 0.01.

**Figure 4.**
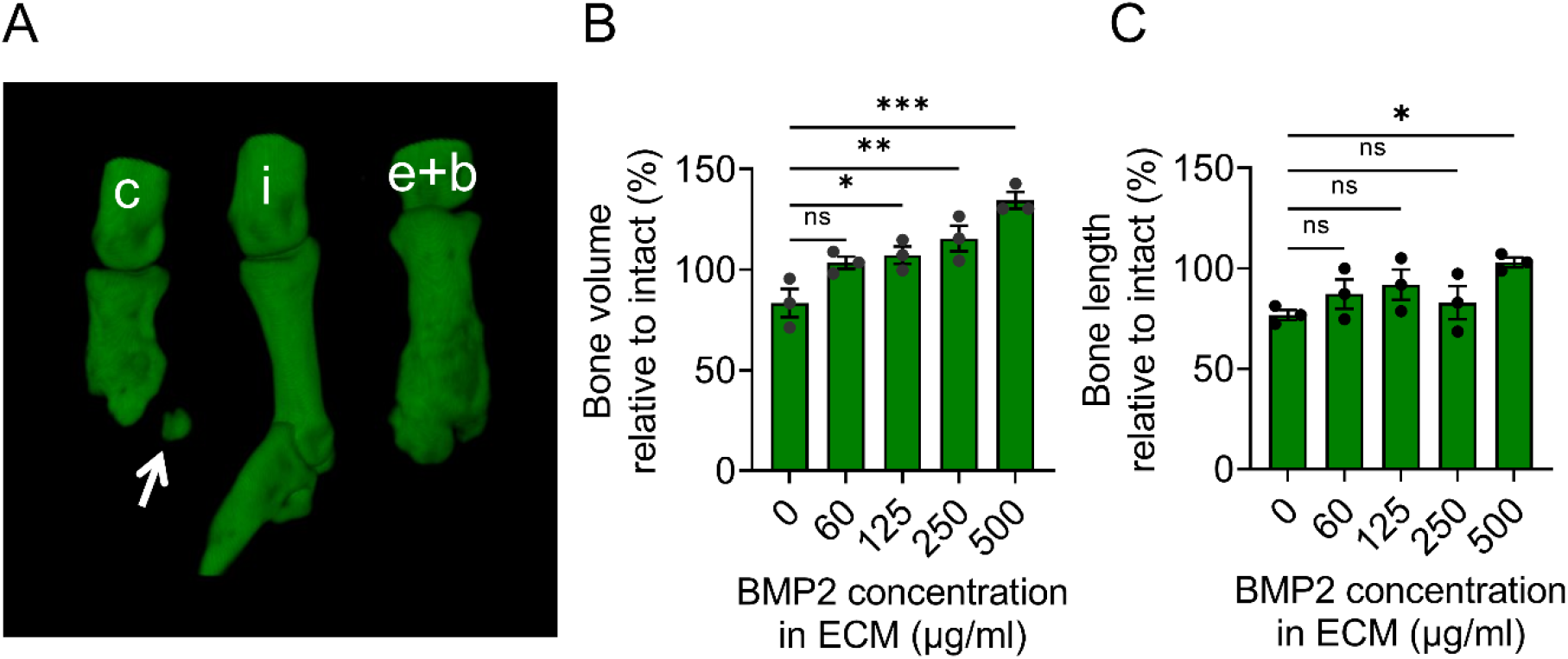
Combination of ECM and BMP2 treatment induced strong bone regeneration of P2 in SPA. (A) micro-CT 3D reconstruction of mice digits of SPA model in 30-35 DPA. c, control group; I, intact group; e+b, ECM+BMP2 group. White arrow indicates sesamoid bone. Top of images indicates proximal site. (B, C) Bone volume and length of mice digit received ECM, BMP2 combinational treatment and intact digit were obtained by micro-CT analysis in SPA model in 30-35 DPA. Bone volumes and lengths of ECM+BMP2 group were normalized to bone volume and length of intact P2 in middle digits of same back foot. n =3, n indicates the number of biological replicates. One-way ANOVA test (Dunnett’s multiple comparisons test) was performed to compare the overall and time point specific effect of ECM and ECM, BMP2 combinational treatment. *p< 0.05, **p< 0.01, ***p< 0.001.

### SPA model and ECM treatment

For SPA, the mice were anesthetized in a prone position, the digits were fixed to the center of the microscope field of view, and the skin was incised until the P2/P3 joint was visible under the microscope. The mouse digits were amputated at the distal 1/3 of P2, and the amputated bone tissue was removed (distal P2 and P3), preserving soft tissues as much as possible. At the same time, ECM particles were implanted at the bone amputation site, closing the wound while wrapping the ECM particle with preserved soft tissue. Forceps were placed against the fat pad until blood coagulated. No other post-operative treatment was performed.

### Digits processing and histology analysis

After treatment, the mice were euthanized for digits collection. No additional re-amputation was performed during the experimental period. The collected mouse digits were fixed in a 4% formaldehyde solution for 24 h and decalcified for 36 h. Tissues were rinsed under running water for 12 h and then dehydrated with gradient alcohol. Dehydrated tissues were treated with n-butanol and paraffin, cut into 4-6 μm slices, and routinely stained with HE or toluidine blue. The n-butanol and paraffin treatment process was as follows: anhydrous ethanol and n-butanol 1:1 mixture for 2 h, n-butanol I for 3 h, n-butanol II for 2 h, low-temperature paraffin for 1.5 h, high-temperature paraffin I for 1.5 h and high-temperature paraffin II for 6 h. The slices were sealed with neutral gum.

### Micro-computed tomography (micro-CT) scans

The collected hindfeet were immersed in 70% alcohol and sent to Guangzhou ZhongkeKaisheng Medical Technology Co. for micro-CT scanning. Images and bone volumes and lengths were processed and obtained using ImageJ software [42] and Python [43].Statistical analysis was performed using Graphpad prism (v9.5.0) software.*p*< 0.05 was considered statistically significant.

## 3. Results

### 2.1 SPA model showed potent bone regeneration

To investigate the effect of ECM particles in the regeneration of mice digits, we established SPA model, where P3 and distal portion of P2 is removed while retaining soft tissue to encapsulate ECM particles adjacent to distal portion of amputated P2 (Fig.1A, B). Due to the methodology in the establishment of SPA model, the appearance of the digits had minor changes compared with intact (Fig.1C). And the ECM and control groups had shown the same digit length of establishment of SPA model(Fig.1C). Alternatively, ECM treated digits showed significantly increased digit diameter compared with control group (Fig.1D, E). We therefore investigated whether bone regeneration is enhanced by ECM treatment in SPA model. We noticed that ECM group had larger amount of new bone in the amputation site compared with control (Fig.1F). Interestingly, the control group in SPA model also exhibited a larger amount of new bone compared with control group in CA model (Fig.1F). These results indicate that bone regeneration is present in SPA model, and ECM treatment may increase the effect of bone regeneration in SPA model.

### 2.2 ECM particle treatment enhanced bone regeneration response in SPA model

To further quantify the regenerative efficacy of ECM particle treatment in SPA model, micro-CT scans were performed on digits at 10, 21, and 35 days post amputation (DPA). New bones appeared on 10 DPA and its amount peaked on 21 DPA. At 35 DPA, new bone density increased to a degree similar to intact bone in control group (Fig.2A). In contrast, ECM particle treated group manifested smaller bone density (manifested by lower micro-CT intensity) compared with control group (Fig.2A). Bone volumes in the ECM group were significantly increased compared with control group (Fig.2A, B). Analysis of osteogenesis indicates that it can be classified endochondral ossification (Fig.S1A, B), which is the same as the osteogenesis CA process described by Dawson et al[20].

Furthermore, the presence of cartilage in the amputation site is observed after 26 DPA in the control and 34 DPA in ECM groups of SPA model (Fig. 2C), indicating that bone regeneration is active on 26 DPA in SPA model, whereas BMP treatment did not induce bone regeneration at 24 DPA in CA model[21]. These results suggest that the ‘regeneration window’ of the SPA model may be longer than the 24 DPA of the CA model.

Additionally, we noticed the presence of free-floating bones in the ECM group but not in the control group (Fig. 2D, E; Fig. S1C, D), This suggests that ECM particle treatment induced the formation of more than one independent free-floating bones in SPA.

### 2.3 Low bone mineral density (BMD) in ECM-induced regenerated bone

ECM particles treatment increased bone volume of mice digits in SPA model, furthermore, we noticed new bones in ECM group tend to have lower bone density compared with control group. To quantitatively validate our observation, we processed the micro-CT greyscale data to obtain BMD values, as reported by Hoffseth et al [43]. BMD values were divided into 3 categories: high (>2.01 g/cm^3^), medium (1.50 ∼2.01 g/cm^3^), and low (0.96 ∼1.49 g/cm^3^), and the proportion of each category was calculated. In intact P2, the largest proportion was the high-density zone, and the smallest proportion was the low-density zone (Fig. 3A). Both the control and ECM groups exhibited the lowest BMD at 10 DPA, with a subsequent increase over time. This resulted in a gradual decrease in the proportion of low density and a corresponding increase in the proportion of high density (Fig. 3A). The percentage of low-density zone in the ECM group was higher compared with the control group, and the percentage of high-density zone was significantly lower compared with the control group (Fig. 3A). This suggests that the amount of chondrogenesis in the ECM group was higher than that in the control group, and the degradation of residual bone in the ECM group was more than that in the control group.

Furthermore, we compared the mean BMD between ECM treated and control group. Both ECM and control group had smaller bone density compared with intact (Fig. 3B). The BMD gradually increased overtime in ECM and control groups, however, BMD in ECM group is significantly reduced compared with control group (Fig.3B). These results indicate that ECM induced bone regeneration is characterized by the presence of low bone density before the 35 DPA.

### 2.4 ECM+BMP2 treatment robustly induced bone regeneration in SPA model

Previous studies indicates that treatment with BMP2 significantly increases the length of P2 bone in CA [38]. Therefore, we investigated whether BMP2 treatment could exert synergistic bone regeneration effects with ECM treatment in SPA model. We adjusted the amputation site proximally to the midway of the P2 bone, creating more space for new bone.

We quantified the bone volume and length by micro-CT assay at 30-35 DPA. We found that combinational treatment of ECM and BMP2 had greater regenerative capacity compared with digits treated by ECM alone (Fig.4A). The increment of bone volume induced by combination treatment is dose dependent on BMP2 concentration (Fig.4B). Interestingly, in all groups receiving ECM+BMP2, bone volume exceeded that P2 of the intact group (Fig.4B). These results indicate that combination treatment robustly induced bone regeneration in SPA model, exceeding the intact group, which we characterize as excessive bone regeneration. And we observed an increased bone length increment by combinational ECM and BMP treatment (Fig.4C).

## 4. Discussion

In the CA model, both bone and soft tissue were removed, whereas in the SPA model, the bone was removed while the soft tissue was retained. This retained soft tissue was originally intended to hold the solid ECM; however, unexpectedly, significant bone regeneration was observed in the SPA model compared to the CA model in the absence of ECM treatment. This result suggests that preservation of soft tissue shifts the healing response after amputation toward a regenerative response. Soft tissues could provide a favorable microenvironment for bone regeneration in SPA model, which is consistent with the view that the root cause of regeneration failure after amputation is not the lack of regenerative response cells, but rather the unfavorable wound microenvironment that limited regenerative potential[11, 16]. For example, Dawson et al. found that blastema was also generated in proximal amputated P3, but the generated blastema failed to complete regeneration [13], indicating the presence of underestimated regenerative potential of mice digits.

In this study, we found that ECM particles treatment significantly induced bone regeneration in mice digits, which promoted the formation of new bone and free-floating bones in SPA model. BMP is known for its strong bone regeneration effect. Injecting BMP into soft tissues like muscle or subcutaneous tissue can create bone, also known as ‘ectopic bone’[44, 45]. Although ECM cannot generate ectopic bone, the generation of free-floating bone in this study is strong evidence that ECM has a bone regenerative effect. The presence of free-floating bone could be associated with the nature of ECM particles treatment: the ECM particles are extruded ECM lyophilized powders, they may disperse due to mechanical stress during ECM implantation. The dispersed ECM fragments can separate form bones, i.e., free-floating bone. And we also noticed that ECM induced bone regeneration is characterized by a low new bone density before 35 DPA. We think that lower new bone density is pro-regenerative as it correlates with an extended period required to achieve normal bone density and enlarged bone extension space. Thus, leads to enhanced bone regeneration.

The location of the inducer is an important factor in determining the shape of the new bone. For instance, the correct placement of gel beads containing BMP2 results in a significant increase in the length of P2, while incorrect placement induces deformation bone [38]. Unlike the gel bead, ECM particles cannot be placed stably in SPA model. During implantation, extruded ECM particles may break and disperse, and when the dispersed ECM are far apart, they generate free-floating bone, while when they are close to each other, they form irregular P2 (Fig.S1 C, D). Therefore, the presence of free-floating bone and irregularly shaped bone in this study is related to the location of the ECM, indicating that quantification of bone volume is an accurate estimation of bone regeneration in this study[46-48].

It is feasible to induce bone lengthening in the human finger with ECM particles as the human finger can provide stable and ample space for the ECM particles, preventing them from breaking under stress. It should be noted that ECM particle treatment should be performed in the ‘regeneration window’, to exert its regenerative effect. Indeed, solid-state ECM treatment did not induce bone regeneration in an already-healed human finger injury [49]. Furthermore, treatment of BMP2 did not induce regeneration of an already-healed P2 bones in mice, but the regenerative response to BMP2 was restored after re-injury [50]. Re-injury of the bones while preserving the soft tissue is an approach of gaining the ‘regeneration window’ and it is an important factor for consideration for the implantation of the ECM[51, 52]. ECM/BMP2 has been licensed for clinical use. Although clinical cases of finger/limb regeneration have not been reported, ECM/ECM+BMP2 is a promising target to induce finger/limb regeneration.

This study also indicates that the ECM may be used therapeutically in injures where soft tissue predominates over bone, ECM or ECM+BMP2 can be implanted into the soft tissue, promoting bone regeneration. Conversely, when bone protrudes from soft tissue, the traditional surgical approach is to remove the exposed bone, allowing bone to be encased within soft tissue to prevent necrosis during healing. However, as ECM has been used clinically in the treatment of soft tissue injuries such as skin and muscle [53, 54], the application of ECM to cover protruding bone has the potential to avoid necrosis, thereby eliminating the necessity of removing exposed bones, which is expected to improve the prognosis.

On the other hand, little is known about how to modulate the shape of regenerated fingers/limbs. For example, the relationship between inducer composition/dose and the amount of regenerated bone/soft tissue, the relationship between inducer topology and the shape of a regenerated bone, and the preferred method of bone re-injury: drilling or amputation? Such studies require larger animal models for further study.

## 5. Conclusion

Our results revealed new bone formation at the distal end of P2 in the SPA model, indicating that—unlike in the CA model—bone can regenerate in the SPA model. The preserved soft tissue not only serves to anchor the solid particles but also stimulates bone regeneration. ECM particles increased bone volume, reduced bone density, and promoted the formation of free-floating bone. These findings indicate that ECM plays a definite role in inducing regeneration of the amputated digit. Furthermore, ECM combined with BMP2 elicited a robust bone regeneration response, with the volume of regenerated bone exceeding that of intact P2 bone. Given that both ECM and BMP have already been approved for clinical use, this study demonstrates that the conditions are now in place to conduct clinical trials on ECM-based regenerative therapy.

## Declarations

### Ethical Statement

The animal experimental protocol was approved by the Experimental Bioethics Committee of Guilin Medical University. Approval number: GLMC202303159.

### Data availability statement

The measurement data and detailed statistical analysis are available as supplementary table.Datasets used and/or analyzed in the current study are available from the corresponding author on reasonable request.

## Acknowledgements

We thank Prof. Ken Muneoka and his team of Texas A&M university, USA.

## Author contributions

**Yingqin Liu**: Writing – review & editing, Writing – original draft, Supervision, Resources, Project administration, Visualization, Data curation, Formal analysis, Conceptualization. **Boyu Li**: Writing – review & editing, Visualization, Validation, Methodology, Formal analysis, Data curation. **Chunmei Bao**: Writing – review & editing, Visualization, Data curation. **Luting Zengi**: Writing – review & editing, Visualization, Data curation. **Zijian Wang**: Writing – review & editing, Visualization, Data curation. **Xin Sun**: Writing – review & editing, Writing – original draft, Software, Formal analysis, Conceptualization. **Guangchen Sun**: Writing – review & editing, Writing – original draft, Resources, Investigation, Supervision, Conceptualization.

