## Supplementary material for "Extracellular matrix particle treatment induces digit regeneration in soft-tissue preserved amputation (SPA)model of adult mice": Suppliemental File 1

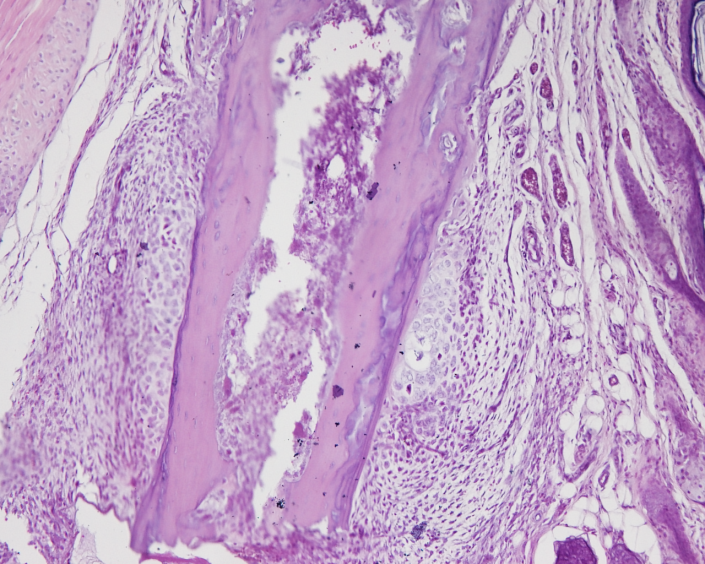

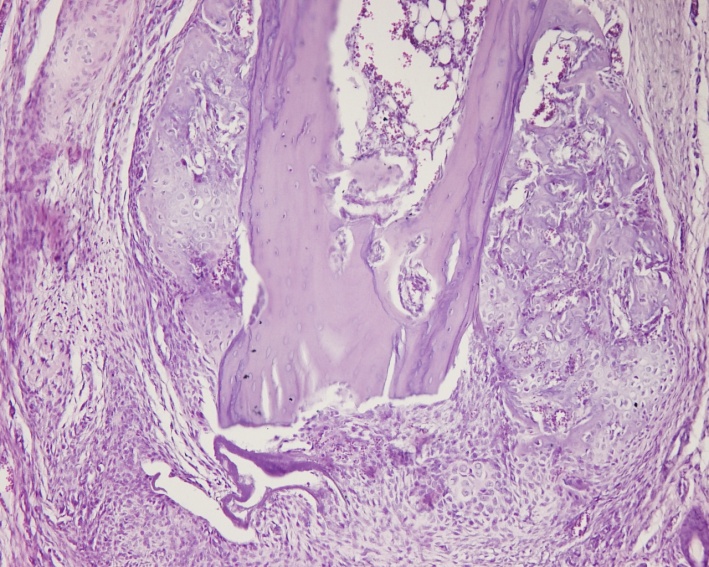

(A). SPA 7d

(B). SPA 14d

b

b

A

B

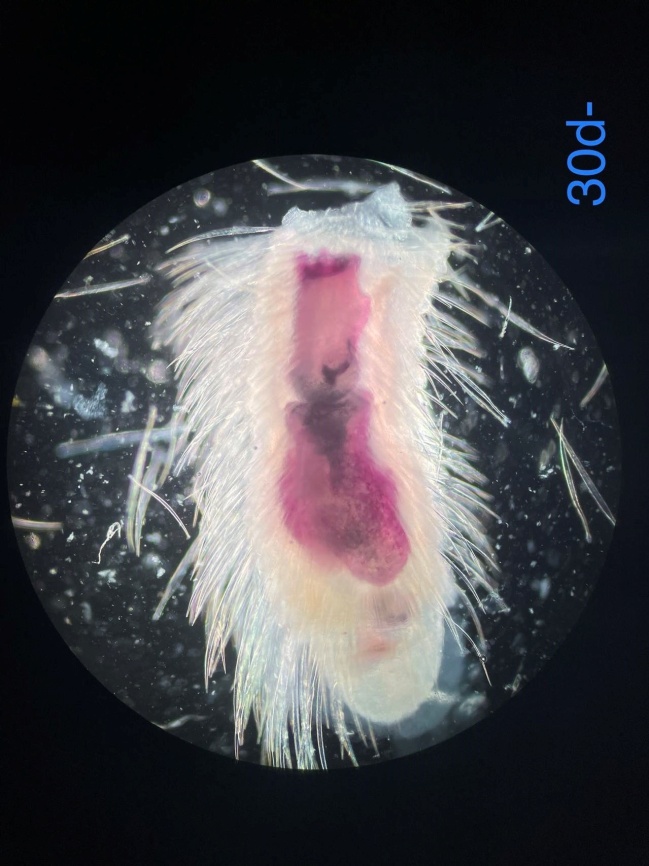

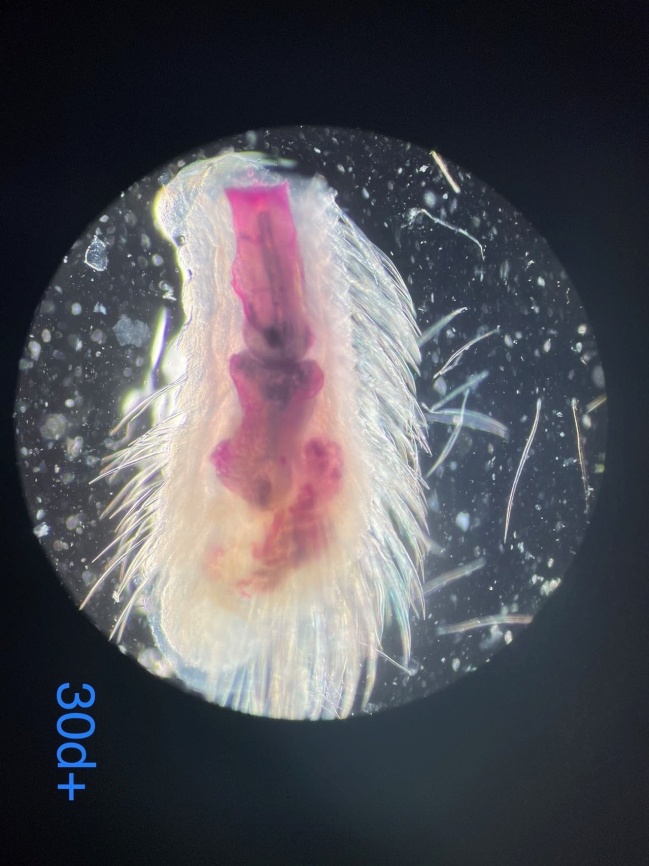

(C). SPA-Control

(D). SPA-ECM

Fig. S1

Endochondral ossification of SPA. (A) Cartilage begins to appear at the periosteum at 7 DPA. The black dotted line is the periosteum of the P2 stump, and cartilage is within the red dotted line. (B) At 14 days, on the P2 side, newly generated cartilage and bone are intertwined. Black dotted line: newly generated bone/cartilage. Red dotted line: cartilage. HE staining of paraffin sections. b, bone tissue. Top is proximal, bottom is distal. scale=200 µm. Figure S2. Free-floating bone/irregular bone in the ECM group in SPA. (A) 30 DPA, no free-floating bone in control group. (B) 30 DPA, the ECM group has free-floating bone (arrow), which may remain free or may fuse with P2, giving P2 an irregular shape. Whole mount bone staining with alizarin red. Top is proximal, bottom is distal.

C

D

Figure.S1. (A-B)Endochondral ossification of SPA. (A) Cartilage begins to appear at the periosteum at 7 DPA. The black dotted line is the periosteum of the P2 stump, and cartilage is within the red dotted line. (B) At 14 days, on the P2 side, newly generated cartilage and bone are intertwined. Black dotted line: newly generated bone/cartilage. Red dotted line: cartilage. HE staining of paraffin sections. b, bone tissue. Top is proximal, bottom is distal. (C-D)Free-floating bone/irregular bone in the ECM group in SPA. (C) 30 DPA, no free-floating bone in control group. (D) 30 DPA, the ECM group has free-floating bone (arrow), which may remain free or may fuse with P2, giving P2 an irregular shape. Whole mount bone staining with alizarin red. Top is proximal, bottom is distal.
